# Optical Tweezer Stretching of Miniature Coarse-Grained Red Blood Cells

**DOI:** 10.1101/2020.10.09.333989

**Authors:** P. Appshaw, A. M. Seddon, S. Hanna

## Abstract

Due to the high computational cost of full-cell coarse-grained molecular dynamics modelling, being able to simulate “miniature” cells that effectively represent their full-sized counterparts would be highly advantageous. To accurately represent the morphological and elastic properties of a human red blood cell *in silico*, such a model is employed utilising the molecular dynamics package LAMMPS. The scale invariance of the model is first tested qualitatively by following the shape evolution of red blood cells of various diameters, then quantitatively by evaluating the membrane shear modulus from simulations of optical tweezer-style stretching. Cells of physical diameter of at least 0.5µm were able to form the characteristic biconcave shape of human red blood cells, though smaller cells instead equilibrated to bowl-shaped stomatocytes. A positive correlation was found between the cell size and both magnitude of deformation from optical tweezer stretching and scaled shear modulus, indicating a lack of scale invariance in the models elastic response. However, the stable morphology and measured shear modulus of the 0.5 − 1.0µm diameter cells are deemed close enough to past *in vitro* studies on human red blood cells for them to still offer valuable use in making simplified predictions of whole-cell mechanics.

**SIGNIFICANCE:** The study tests the invariance of a coarse-grained molecular dynamics red blood cell (RBC) model to system scale, asking whether it is qualitatively and quantitatively viable to perform whole-cell simulations in “miniature”. Simulating cells at a reduced scale greatly improves computational speed, making possible computational experiments that would otherwise be too computationally demanding. This facilitates the simulation of larger systems, both in number of whole-cells, and cells of greater structural complexity than the RBC. More generally, the accurate and efficient modelling of biological cells allows computational experimentation of real-world systems that would be very challenging or impossible to perform *in vitro*. Therefore, miniature-cell modelling could help both direct development in whole-cell modelling, and also developments in more widespread bio-physical studies.

## 1 INTRODUCTION

The red blood cell (RBC) is the simplest and most well researched blood-borne cell, making it an ideal candidate on which to develop techniques in whole-cell computational modelling (1–3). The RBC is a highly deformable, “rubbery” cell, able to recover its shape after squeezing through very narrow capillaries (4). The cell is primarily comprised of a 2-component membrane, enclosing a cytoplasm fluid interior. Due to the entirely viscous nature of the cytoplasm, the cell membrane is solely responsible for the elastic response of the cell (5). As the membrane thickness is much lower than the diameter of the whole cell, it has 3D structure describable by 2D elastic parameters (6). The resistance to extension of the whole-cell is then characterised by two properties of the membrane: the out-of-plane bending rigidity *B*, and the in-plane shear modulus *µ*_*s*_ (7). The RBC membrane is composed of a lipid bilayer and distinct cytoskeleton network, connected by transmembrane proteins. The lipid bilayer is essentially a 2D fluid-like structure embedded in 3D space, resistant to bending, but unable to sustain in-plane shear stress due to its highly diffusive nature (8). Conversely, the cytoskeleton provides the resistance of the membrane to shear deformation, being a structural network bound to the inner surface of the bilayer. Therefore, the bending rigidity *B* of the cell membrane is dominated by the lipid bilayer, whereas the shear modulus *µ*_*s*_ is dominated by the cytoskeleton (8–10).

Many different formalisms have been used to develop computational models of RBCs. To date, the most popular have been continuum methods such as the finite element method (FEM), and the particle-based dissipative-particle-dynamics (DPD) approach (2, 3, 11). However, recently coarse-grained molecular dynamics (CGMD) has seen increasing popularity (8, 12–15). In CGMD, an atomic system is coarse-grained (CG) into a more computationally efficient representation, with CG particle interactions managed by inter-particle potentials. The length-scale of the system is defined by a specific parameter of the system, such as the thickness of the bilayer. The elastic properties are then dictated by the broader physics of the complete system and not explicitly specified by the interaction potentials. Conversely, comparative DPD and continuum models require the *a priori* knowledge of elastic moduli, made explicit within the model functions (15, 16). Therefore, CGMD is unique in allowing evolution of these properties, crucially enabling the testing of how elastic parameters change under new stimuli.

However, CGMD is very computationally expensive, with membrane models historically restricted to simulation of only small patches of membrane (8, 17). Only recently has a full-scale CGMD RBC been modelled in its entirety (15). Despite being an implicit-fluid model with notable care taken to ensure computational efficiency, simulating the full-scale cell (consisting of 3.2 million particles) for 100, 000 time-steps on 20 CPU cores still took on the order of a day. An alternative speed-up approach has been to simulate “miniature” cells, consisting of far fewer particles, but whilst adopting the same length scaling (14, 18). These cells thus have a significantly reduced physical diameter, but assume elastic properties equivalently representative to those of a full-scale cell. To our knowledge however, no quantitative verification of this “miniature cell” approach has been conducted.

To date, most CG particle models of RBCs have only been validated qualitatively, by studying their shape at rest and under flow conditions (19). To determine the membrane shear modulus, experimental techniques typically involve applying some external force in order to deform it. Historically, most quantitative data on the RBC shear modulus has come from the technique of micropipette aspiration (20). By drawing a protrusion of the membrane into a pipette, the relationship between applied pressure and protrusion extension allows approximation of *µ*_*s*_, creating an accepted range for the RBC membrane of 4 − 10µN/m (21). Recently however, optical tweezers have seen increasing popularity as an alternative (12, 21–23). Optical tweezers make it possible to apply pN-level forces to a membrane, either directly or through manipulation of attached microbeads (5). In the latter case, two silica microbeads attached at opposite ends of a cell membrane can be used like handles to stretch the cell. Analogously to the micropipette technique, the gradient of the resulting force-extension curve then allows approximation of *µ*_*s*_ (5, 7). Analysis of deformation responses from the optical tweezers technique has shown a wide range in calculated RBC membrane shear modulus, with Henon *et al*. finding 2.5µN/m, Mills *et al*. 5.3 − 11.3µN/m, and Sleep *et al*. up to 200µN/m (5, 7, 21).

This work gives, to the best of our knowledge, the first quantitative assessment of the “miniature cell” approach to CGMD whole-cell modelling, by testing the scale invariability of a fundamental elastic parameter of a human RBC; the membrane shear modulus *µ*_*s*_. Firstly, the simulation methodology is outlined, before attempting the evolution of initially spherical cells of various model diameters to a biconcave stable state. The evolved whole-cell systems are then used as input for simulated optical tweezer stretching tests, with *µ*_*s*_ determined via a dimensional scaling. These results are then compared to past *in vitro* studies, and used to discuss the invariability of aspects of the model to the physical cell diameter.

## 2 MATERIALS AND METHODS

The model used in this work follows closely that of Fu *et al*. (14), who built upon the lipid membrane model of Yuan *et al*. (24). The model is that for a CGMD, 2-component RBC, with explicit representations of the lipid bilayer, cytoskeleton, and internal and external fluid particles (see Figure 1A). The lipid bilayer is represented as a one-particle-thick monolayer of CG spherical particles, each representing a large number of constituent lipids. In determining a distance dependant function for lipid-lipid interactions in the bilayer, it is challenging to find a form that produces the correct diffusion of particles. It has been shown that the classical 12-6 Lennard-Jones (LJ) potential only produces two membrane phases, a solid at low temperatures and gas at high temperatures (25). At small separations the inter-particle forces are too strong to permit particle diffusion, and at large separations too weak to keep particles bound together. To provide the intermediate fluid phase necessary to allow such behaviour, a two branch interparticle function can be adopted (24).

**Figure 1:**
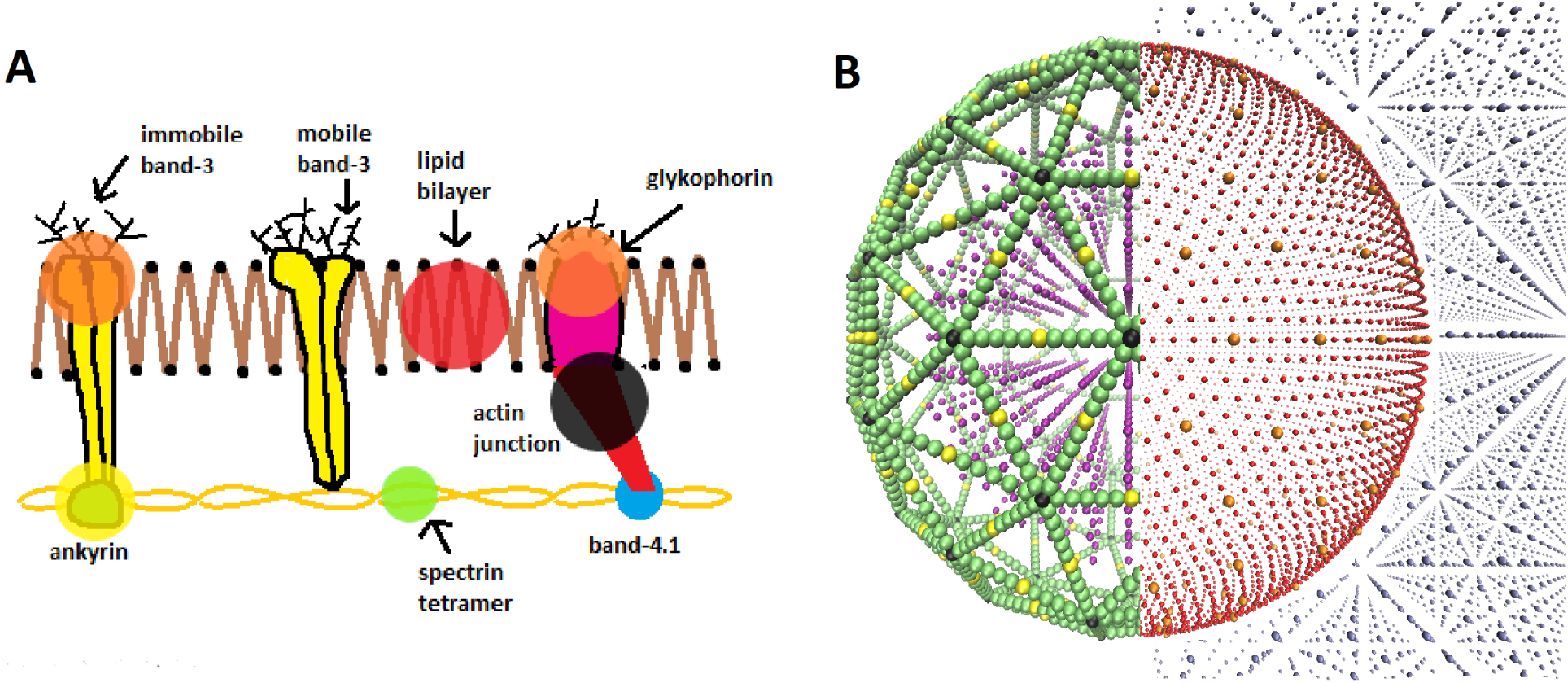
**(A)** Pictorial representation of the RBC membrane components, showing the cytoskeleton network attached to the lipid bilayer. Actin junction complexes (actin protofilament and protein band-4.1) connect the spectrin tetramers. The cytoskeleton is tethered to the lipid bilayer via transmembrane proteins - immobile band-3 at the spectrin-ankyrin binding sites and glycophorin at the actin junctional complexes. **(B)** Graphic of our pre-evolved (spherical) RBC, with each CG particle type shown in colour: (red) lipids, (orange) trans-membrane proteins, (green) spectrin tetramers, (black) junction complexes, (yellow) ankyrin, (purple) internal fluid and (grey) external fluid. The graphic is split into two mirror halves, to make the distinct particle types clearer visually. The particles described in Figure **B** are highlighted in Figure **A** by opaque circles of corresponding colour.

This work adopts the lipid-lipid interaction potential of Yuan *et al*. (24), hereafter referred to as the Yuan potential. The Yuan potential has been shown to represent well the mechanical properties of a RBC bilayer membrane, including a diffusive fluid phase, due to the separation of attractive and repulsive branches (14). It has an orientational dependence which allows the complex lipid hydrophobicity to be represented, being essential for the self-assembly of the bilayer in an aqueous environment (8, 26). The membrane properties of spontaneous curvature *c*_0_, bending rigidity *B* and diffusivity *D* are conveniently characterised by three Yuan model parameters, *θ*_0_, *µ*_*Y*_ and *ζ* respectively. *θ*_0_ signifies the most energetically favourable angular configuration between particles, with *µ*_*Y*_ weighting the energy penalty for deviation away from this. *ζ* controls the slope of the attractive branch of the potential. The potential also features the LJ-like parameters of length *σ*, energy well depth *ϵ* and cut-off radius *r*_*c*_. See the Supplementary Material S1 for further detail on the formalism of the potential.

While bilayer-bilayer interactions are managed by the Yuan potential, all other particle-particle interactions operate through classic 12-6 LJ potential functions. Unless otherwise stated, interactions operate under parameters following Fu *et al*. (14), summarised in Table 1. Assuming a typical RBC curvature *c*_0_ ∼ −0.5µm^−1^, the membrane curvature is parameterised as sin *θ*_0_ = − 1.41 × 10^−3^ (27, 28) (see the Supplementary Material S1 for detail on this calculation). A Hookean harmonic bond-potential is also used to couple the transmembrane proteins of the lipid bilayer to the cytoskeleton. The characteristic biconcave RBC shape can only be attained if the spectrin network undergoes constant structural remodelling, so as to relax the in-plane shear elastic energy to zero (8). One way to ensure the cytoskeleton is initially stress free is to modify the equilibrium bond length in the bond-potential so that the bond energy is initially zero. This is achieved by making the equilibrium bond length a variable corresponding to the initial bond lengths between each particle pair, rather than a constant as in the standard Hookean potential (14).

**Table 1:** Default pair potential parameters for each CG particle interaction type within the model, all given in LJ units. The mass of each CG particle type is also given. ‘Fluid’ refers to both internal and external solvent. ‘Bilayer’ refers to the lipids and transmembrane proteins. ‘Cytoskeleton’ refers to spectrin, junction complex and ankyrin particles. Values that differ from the Fu *et al*. (14) implementation are marked with a *.

| Particle Interaction | Pair Potential | $\sigma$ | $\epsilon$ | $r_{cut}$ | $\zeta$ | $\mu_Y$ | $\sin \theta_0$ |
| --- | --- | --- | --- | --- | --- | --- | --- |
| fluid-fluid | LJ | 2.7 | 0.2 |  |  |  |  |
| fluid-cytoskeleton | LJ | 1.0 | 0.2 |  |  |  |  |
| cytoskeleton-cytoskeleton | LJ | 1.0 | 0.2 |  |  |  |  |
| bilayer-bilayer | Yuan | 1.0 | 1.0 | 2.6 | 4 | 3 | $-1.41 \times 10^{-3*}$ |
| bilayer-cytoskeleton | LJ | 1.0 | 0.2 |  |  |  |  |
| bilayer-fluid | LJ | 1.0 | 0.2 |  |  |  |  |
| **Particle Type** | $m^M$ | | | | | | |
| bilayer, cytoskeleton | 0.5* |  |  |  |  |  |  |
| fluid | 1.0* |  |  |  |  |  |  |

Simulations are run utilising the LAMMPS molecular dynamics coding package (29), operated as a library within Python. LAMMPS handles the thermodynamic evolution of the system, while particular biophysical calculations are performed in the parent Python code. Initial particle configurations are input from an independent Python code which generates a 3D “supercell” volume containing the configuration of pseudo-particle types as required (see Figure 1B), alongside particle classifications interpretable by LAMMPS. Simulations are performed in the system of non-dimensionalised LJ units. However, to compare results with past *in vitro* studies, quantities must then be converted to SI units. In the presented formalism, each variable *i* is associated with a dimensional conversion parameter *σ*_*i*_ which relates the non-dimensional “model” LJ unit (denoted *i*^*M*^) to “real” SI unit (denoted *i*^*R*^). For example, the conversion of a length *r* from LJ units to meters is denoted *r*^*R*^[m] = *σ*_*r*_ [m]*r*^*M*^. The unit conversions used in this work are length-scale *σ*_*r*_ = 5nm, time-scale *σ*_*t*_ = 80ns, energy-scale *σ*_*ϵ*_ = 1.8 × 10^−20^J, force-scale *σ*_*F*_ = 3.5pN, pressure-scale *σ*_*P*_ = 1.4 × 10^5^N/m^2^, temperature-scale *σ*_*T*_ = 1.3 × 10^2^K and mass-scale *σ*_*m*_ = 4.9 × 10^−6^kg. All simulations were run on 1 − 4 nodes of the local supercomputer, with each node having two 14 core 2.4 GHz Intel E5-2680 v4 (Broadwell) CPUs, and 128 GB of RAM. See the Supplementary Material S2 and S4 for detail on the conversion of each unit and benchmark of the model respectively.

Implementation of the Yuan potential is verified against Fu *et al*. (14) by testing of the bending rigidity and diffusivity from the thermal fluctuations of a patch of isolated bilayer membrane. Fu *et al*. sets the mass of the bilayer particles to 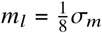 and all other particles to 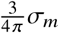. However, we found that in our implementation of the model this failed to reproduce comparative elastic and diffusive properties. Instead, we found better agreement by setting the mass of all membrane particles to 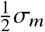, and all other particles to *σ*_*m*_. This gave a rigidity of *B* = 18.3*k*_*B*_*T*, matching well with the result of *B* = 18*k*_*B*_*T* by Fu *et al*. (14). Similarly, an approximate simulation time scale of *σ*_*t*_ = 80ns was then obtained from the diffusivity in the bilayer, matching closely with the value *σ*_*t*_ = 100ns of Fu *et al*. See the Supplementary Material S3 for detail regarding these tests.

## 3 RESULTS

### Cell Equilibration

To test the miniature-cell methodology, whole-cells are generated at various model diameter *D*^*M*^, with smaller cells comprising proportionally fewer CG particles. Each cell is generated as an initially spherical configuration of CG particles suspended within a fluid of water-like CG particles. The cell membrane is generated as two concentric spherical shells of bilayer and inner cytoskeleton, enclosing an internal CG fluid representing the cytoplasm. Each particle type is then thermodynamically activated sequentially as follows:

- An isothermal-isobaric (NPT) ensemble is applied to the external water, and the canonical ensemble (NVT) to the internal fluid over 25,000 steps.
- The spectrin, ankyrin and junctional complexes of the cytoskeleton are then equilibrated with the NPT ensemble over 25,000 steps.
- The lipids and transmembrane proteins in the bilayer are the last to be equilibrated, using the NVT ensemble over 50,000 steps.

Each cell is equilibrated from its initially spherical state with time-step length *τ* = 0.02*σ*_*t*_, pressure *P*^*M*^ = 0.05, and temperature ramping from *T*^*M*^ = 0.02 to *T*^*M*^ = 0.23 (14). For these initial shape-evolution simulations, some model parameters are changed from their default values: the membrane particles have their mass increased to *σ*_*m*_, and the energy well depth in the Yuan potential is increased to *ϵ* = 1.5, to strengthen the lipid bindings, temporarily.

Membrane folding is then induced by reducing the initial number of internal fluid particles *N*_*IF*,0_ to a final number *N*_*IF*_ (see Table 2). Small, equal and opposite forces are also applied to circular areas on each *XY* face of the cell to promote biconcave indents to manifest perpendicular to the *XY*-plane. Particles are deleted gradually to a final fraction of internal fluid particles *n*_*IF*_ = *N*_*IF*_ */N*_*IF*,0_, differing with cell size. The rate of compression has an effect on both the equilibrium shape and stability of the transition (6, 14). All cells have *N*_*IF*_ reduced at a constant rate of 3% every 5000 time-steps, which was determined to be most conducive to achieving a biconcave final state. The concave regions of the cells then develop gradually with the compression. Deviation from this rate results in alternative unwanted vesicle transitions such as to prolate, dumbbell rods, or inward or outward budding (see Yuan *et al*. for examples (6)).

**Table 2:** Equilibrated final states of each RBC cell size, with final ratio *V/R*^3^ achieved from chosen compression fraction *n*_*IF*_. The number of bilayer to total particles is also given, as-well as the number of steps run in the compression stage of the simulation.

| Final State |  |  |  |  |  |
| --- | --- | --- | --- | --- | --- |
| $D^M$ | 50 | 75 | 100 | 125 | 150 |
| $V/R^3$ | 2.02 | 1.90 | 2.08 | 2.03 | 2.00 |
| $n_{IF}$ | 0.68 | 0.49 | 0.42 | 0.37 | 0.32 |
| Bilayer particles | 8,346 | 18,704 | 33,400 | 52,069 | 75,156 |
| Total particles | 34,944 | 114,592 | 269,296 | 521,437 | 896,703 |
| Compression steps | 55,000 | 85,000 | 95,000 | 105,000 | 115,000 |

Only cells with diameter *D*^*M*^ ≥ 100 were able to achieve a biconcave discocyte final state, with smaller cells instead relaxing to bowl-shaped stomatocytes (see Table 2). The degree of biconcavity in a healthy RBC can be characterised by the volume-radius ratio *V/R*^3^ = 1.57 (28). An optimal particle fraction *n*_*IF*_ for each cell is found by slowly deleting internal fluid particles until a target volume-radius ratio is reached. *n*_*IF*_ is found to be inversely proportional to *D*^*M*^ which achieves a consistent volume-radius ratio between cell sizes. Furthermore, below a critical ratio *V/R*^3^ ≲ 1.9 the internal fluid becomes unable to fill the region between the two enclosing membrane edges, closing the gap. This critical ratio does not appear to have a direct relationship with cell-size. However, the effect is more pronounced for the biconcave cells, and thus suppressed in the bowl-shaped *D*^*M*^ < 100 cells. To maintain consistency between cells, *n*_*IF*_ is chosen such that cells are compressed to a ratio *V/R*^3^ = 2.0 ± 0.1, slightly higher than that of a healthy RBC (see Table 2). However, qualitatively, the *D*^*M*^ ≥ 100 cells relax to morphologies closely resembling those of a healthy human RBC (see Table 2).

### Optical Tweezers

Once each cell has been equilibrated to its stable state, it can be used as input for subsequent simulations of optical tweezer stretching. The Yuan potential and membrane particle masses are now set back to their defaults, *ϵ* = 1.0 and 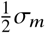 respectively. Stretching is applied by assuming the influence of two spherical microbeads, each of radius 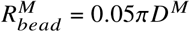 (equivalent to 2.5µm diameter at a full-scale RBC), centred on those lipid particles having the minimum *x*_−_ and maximum *x*_+_ coordinates (see Figure 2A). All the lipids within 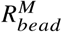 of these extremities are designated “bead” particles. Stretching forces 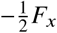 and 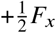 are then applied, split evenly across only these *N*_−_ and *N*_+_ “bead” lipids respectively.

**Figure 2:**
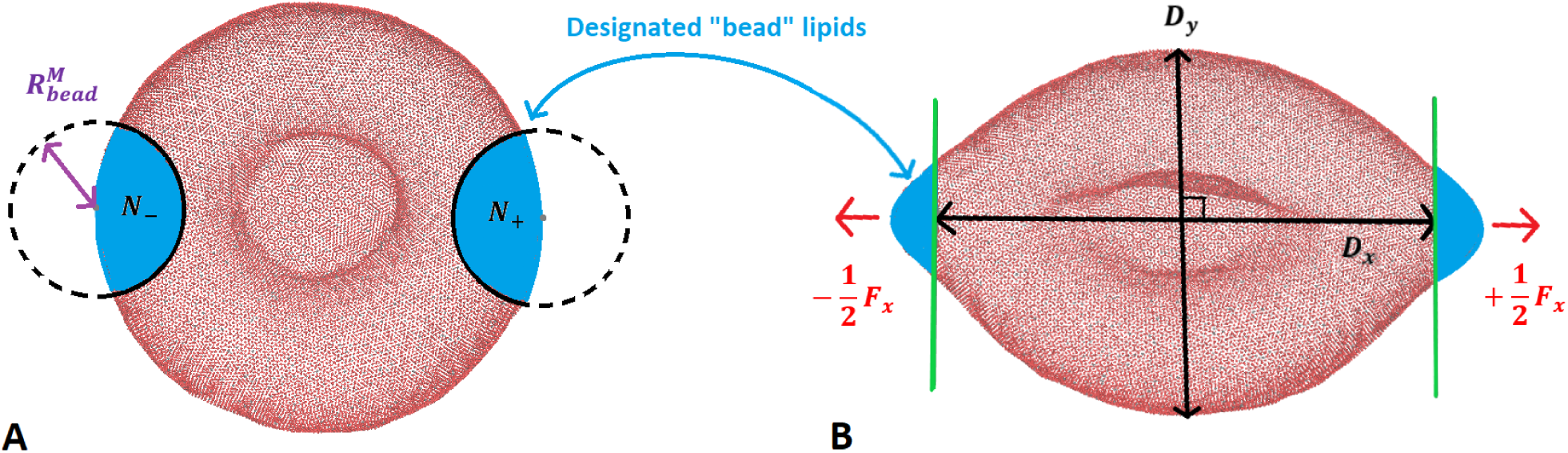
(**A**) Graphic showing the *XY* -plane projection of a cell at initial biconcave equilibrium, with contact areas of each bead designating *N*_±_ lipid particles as “bead” particles. (**B**) Graphic showing the cell after stretching has been applied by forces 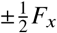, where each force is split evenly across only those designated *N*_±_ bead-particles respectively. The axial *D*_*x*_ and transverse *D*_*y*_ diameters are also shown, with intersecting (green) lines indicating the contact planes from the beads.

As an increasing force is applied, the membrane will stretch along the axial diameter *D*_*x*_ and contract along the transverse diameter *D*_*y*_ (see Figure 2B). The resulting force-extension profiles can then be compared against the literature, and *µ*_*s*_ can be approximated in the low deformation regime (5). Following the recommendation of Siguenza *et al*. (23), *D*_*x*_ is taken as the distance between interior planes intersecting the cell at the inner edge of each bead (see Figure 2B). *D*_*y*_ is then the distance between those lipids with the minimum and maximum *y* coordinates in the plane perpendicular to *D*_*x*_.

Following Henon *et al*. (5), assuming linear elastic theory - valid in the low deformation region - applying equal and opposite forces 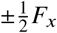 at diametrically-opposed regions of the RBC surface will incur a change in transverse diameter

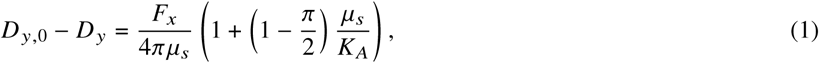

with *D*_*y*,0_ being the stable transverse diameter before stretching. Given that *µ*_*s*_ ≪ *K*_*A*_ for a RBC membrane (6), this expression can be reduced to

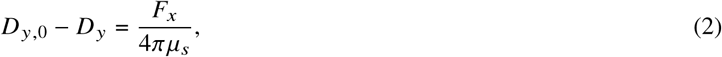

which has been shown to be approximately equivalent for discotic and spherical cells (5).

To represent each cell as a full-size RBC, additional scaling factors are introduced. A constant force-per-particle *f*_*x*_ = 2*F*_*x*_/(*N*_+_ + *N*_−_) is maintained with *D*^*M*^ by including a factor 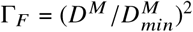 in the total applied force, relative to the diameter 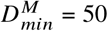 that the model was originally parameterised for. Additionally, a length scaling 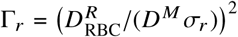 is applied to Eq. 2, relative to the full-scale RBC diameter 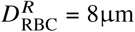. Eq. 2 is thus applied as

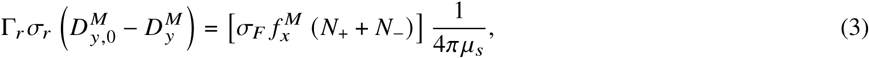

including conversion to SI units.

Plots of extension against time at given force are generated for each cell up to a maximum 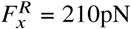, with axial and transverse extension ratios denoted by *λ*_*x*_ and *λ*_*y*_ respectively (see Figure 3A for an example). The force is applied in a step-wise progression from *F*_*x*_ = 0, alternating every 5 × 10^5^ steps between periods of steady ramped increase, and being held constant to allow the cell to stabilise. Each stage of force ramping applies an additional 5*σ*_*F*_ over 5 × 10^5^ steps, corresponding to a rate of 440pN/s. Reduced axial and transverse force-extension curves are then produced (see Figure 3B), where the extension at given *F*_*x*_ is taken as the mean over the relevant stabilisation phase. The smallest (*D*^*M*^ = 50) cell was unable to reach the maximum force of 210pN due to critical membrane rupture occurring above 185pN. The largest (*D*^*M*^ = 150) cell was unable to achieve a maximum force above 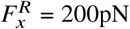, due to computational limitations regarding simulation time and memory access at this system size.

**Figure 3:**
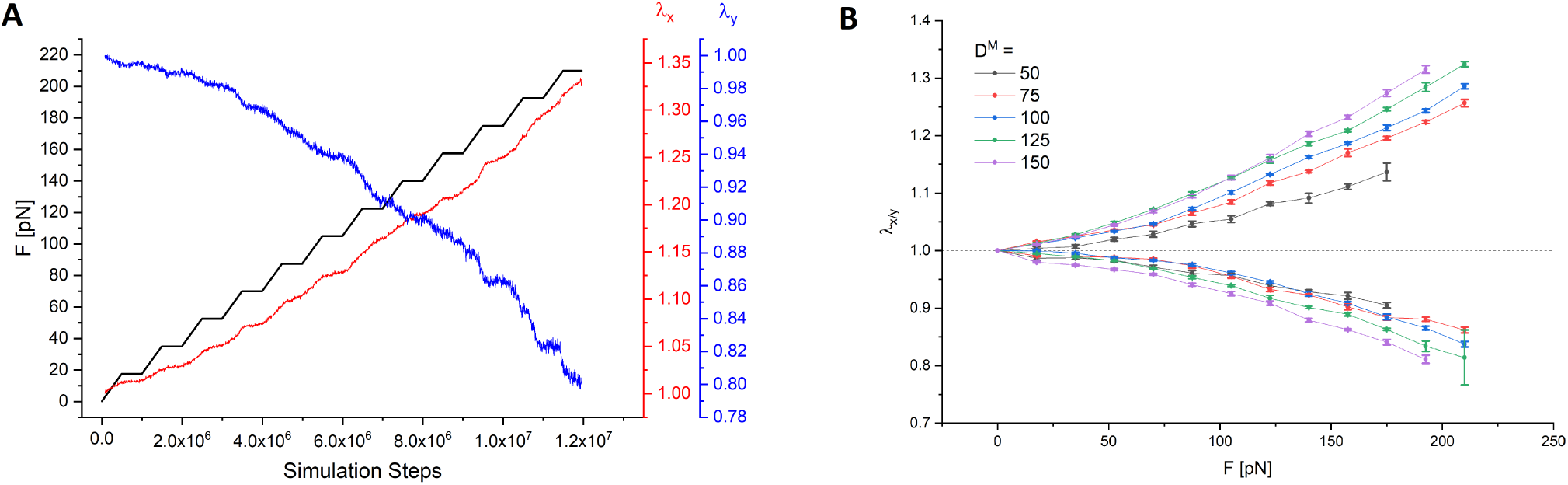
(**A**) Example plot of force and extension against time up to 210pN from the *D*^*M*^ = 125 cell, showing both the axial (red) and transverse (blue) deformation response. The plots from the other cell sizes are given in the Supplementary Material S5. *F*_*x*_ is increased gradually in a step-wise fashion, alternating between ramp and stabilisation stages every 5 × 10^5^ steps. (**B**) Plot showing the combined axial _*x*_ and transverse _*y*_ force-extension curves for each *D*^*M*^ tested. The extension at given force is taken as the mean of the respective stabilisation stage, with error given by the standard deviation in that mean. The *D*^*M*^ = 125 cell has a particularly large error at 210pN, due to having ruptured before the end of that stabilisation phase.

Over the force regime tested, all cell sizes show reduced deformation at given force compared with *in vitro* studies on healthy human RBCs. The experiments of Mills *et al*. (21) saw peak extensions of *λ*_*x*_ = 2.0 ± 0.3 and *λ*_*y*_ = 0.5 ± 0.2 at maximum force of 193pN. Similarly, the experiments of Suresh *et al*. (30) saw *λ*_*x*_ = 2.1 ± 0.2 and *λ*_*y*_ = 0.6 ± 0.1 at 193pN. At the same force, our simulations have resulted in maximum extensions of *λ*_*x*_ = 1.32 ± 0.01 and *λ*_*y*_ = 0.81 ± 0.01 (being those from our *D*^*M*^ = 150 cell). In this high force regime, we have thus found extension roughly three times lower than in these comparative studies (21, 30). The disparity is even greater in the low force regime, where at 39pN our maximum axial and transverse extension are roughly ten times lower comparative to the Mills *et al*. experiment (21).

Regarding the shape of the extension-time curves, observations can be made on the smoothness of the curves, and how continuous the trajectories are. All cells have extension curves follow the step-wise progression of the force, with each period of force ramping producing an increased rate of extension, followed by a gradual flattening of the extension over the subsequent stabilisation phase (see Figure 3A and Figure S4). However, the overall rate of change of extension increases with force above the low deformation regime. This observation is also represented in the reduced force-extension curves (see Figure 3B). The rate of change of extension is initially approximately linear ≲ 80pN, but then gradually increases with growing *F*_*x*_. As *µ*_*s*_ is determined from the gradient of the force-extension curve, this behaviour is consistent with previous studies showing *µ*_*s*_ to be variable over a large deformation regime (21, 31).

To calculate *µ*_*s*_, cells are instead stretched from *F*_*x*_ = 0 up to 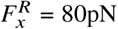 at the same rate of 440pN/s, but now without stabilisation phases. *µ*_*s*_ is determined using Eq. 3 from the gradient within the low deformation limit of the resulting force-extension curves (see Figure 4A). The low deformation regime is defined by the initial region within which the force-extension response remains linear, deemed to be that up to 40pN. Above this region, the rate of extension begins to increase, and linear elastic theory no longer holds, invalidating Eq. 3 (5). The axial and transverse diameters of all cell sizes are seen to randomly fluctuate with time. RBC membranes are known to show shape fluctuation due to thermal noise, introducing an uncertainty in *λ*_*x/y*_ (2). Such random fluctuations are seen to be increasingly prominent at small *D*^*M*^, with much greater noise in the extension (see Figure 4A).

**Figure 4:**
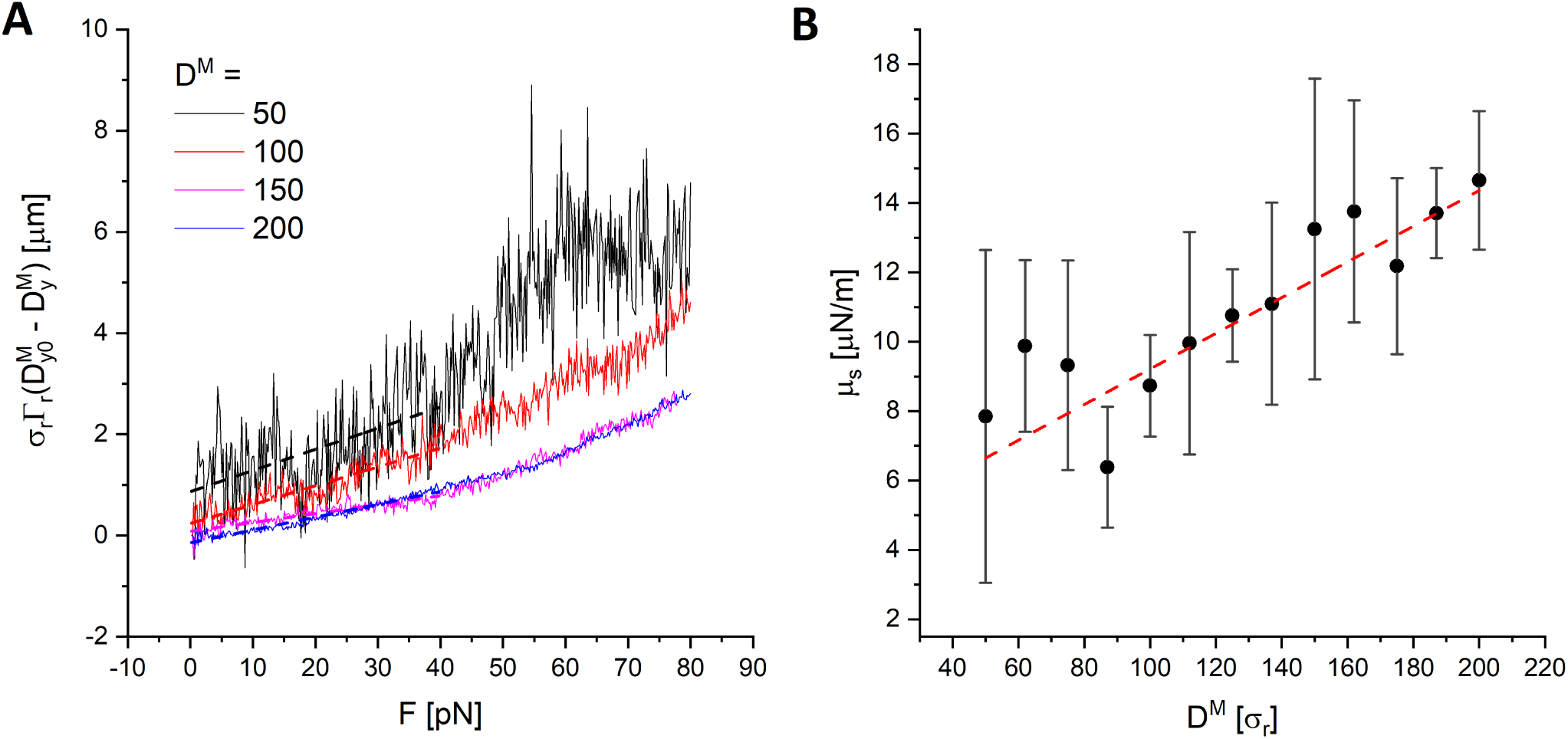
(**A**) Change in scaled transverse diameter against force for the *D*^*M*^ = 50, 100, 150, 200 cells. The shear modulus is then calculated according to Eq. 3 from the gradient in the low deformation limit, with dashed lines of best fit of corresponding colour. (**B**) Evaluated scaled shear modulus against simulated cell size, with dashed line of best fit weighted by error. Standard error is obtained from repeating each simulation 3 times.

Despite dimensional scaling, there appears to be a positive correlation between *µ*_*s*_ and *D*^*M*^ (see Figure 4B). The range *µ*_*s*_ = 6 − 10µN/m evaluated for our *D*^*M*^ < 125 cells is within the accepted range of 4 − 10µN/m from *in vitro* micropipette experiments on healthy human RBCs (21), while our *D*^*M*^ ≥ 125 cells exceed this range (having *µ*_*s*_ = 11 − 15µN/m). However, previous *in vitro* studies using optical tweezer techniques have produced a large range in values for the human RBC shear modulus (*µ*_*s*_ ∼ 2.5 − 200µN/m) (5, 7, 21). Therefore, we do not consider the higher shear modulii of our *D*^*M*^ ≥ 125 cells to be unreasonable for representation of a healthy RBC.

Comparing the reduced force-extension curves between cell sizes (see Figure 3B), we observe a distinct proportionality with *D*^*M*^ - larger cells have a greater deformation response than smaller cells when subject to the same force. If this trend were to persist in our model up to a full-scale cell (that at *D*^*M*^ = 1600), it could provide an explanation for the much lower deformation response seen across our miniature cells compared to *in vitro* studies. Similarly, a positive trend is found between *µ*_*s*_ and *D*^*M*^ from evaluation in the low deformation limit (see Figure 4B). If this trend were to persist linearly, a full-scale cell would expect *µ*_*s*_ ∼ 90µN/m, roughly an order of magnitude higher than the smallest cell tested. However, it should be noted that due to computational limitations from poor model scaling (see Supplementary Material S4), it was infeasible to test cells of *D*^*M*^ ≳ 200. Therefore, it is unclear whether these trends persist beyond the cell sizes tested, or indicate the start of a plateau in the deformation response and measured shear modulus with *D*^*M*^.

## 4 DISCUSSION

In the present computational study, we have observed variations with cell size in the shape of the stable state, shape of the force-extension curve from optical tweezer-style stretching, and shear modulus as measured in the low-force regime. Only cells of *D*^*M*^ ≥ 100 evolved to biconcave discocytes, with smaller cells relaxing to bowl-shaped stomatocytes (see Table 2). However, this is conceptually reasonable, due to the CG nature of the model. By considering all model sizes representative of a full-sized RBC, *D*^*M*^ essentially represents a degree of coarse-graining, as smaller cells are comprised of fewer particles. The more CG a cell, the less versatile it is to shape transitions, due to the reduced number of degrees of freedom. Therefore, it is not unexpected to have found a lower *D*^*M*^ limit on the cells able to form the more complex biconcave shape, nor a complex relationship between *D*^*M*^ and optimal internal fluid fraction *n*_*IF*_.

The effect of cell shape on the stretching response was also investigated. Not only did cells evolve to different total morphology (biconcave discocyte or stomatocyte), cells of the same size would also develop varied non-axial deformities within these shapes. For example, cells would develop as non-axial discocytes where the cell thickness was non-uniform, or having concave regions of varying depth and shape. Furthermore, the orientation of the evolved cell was not always ideal, with concave regions not always manifesting directly perpendicular to the *XY* -face. These effects were particularly prominent in the smaller cells and contributed to inconsistencies in the progression of the extension curves, where reorientation or reconfiguration of the cell shape would produce abrupt changes in extension (see Figure S4). Crucially, these factors often resulted in stretching being initiated on cells having non-axially symmetric *XY* cross-sections, where *D*_*x*,0_ ≠ *D*_*y*,0_. However, stretching tests of cells with *D*^*M*^ = 75 and *D*^*M*^ = 125 and at a variety of initial shapes showed no consistent relationship between the total deformation response and initial shape, in either case. This is consistent with a previous study, which found no significant effect of initial cell shape on the evolution of *D*_*x*_ or *D*_*y*_ (23).

Over the deformation regime tested, all cell sizes showed a reduced extension for a given force compared with previous studies (21, 30). This implies our model membrane to be stiffer than that of the RBCs tested in these studies. A notable consideration to this is the effect of the model bending rigidity on the deformation response. While the Henon analysis implicitly assumes the membrane bending rigidity to have a negligible influence on the deformation response, it has been shown that for a given force, *λ*_*x*_ ∝ *B*^−1/3^, compared to 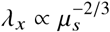 from the shear modulus (32). Following Fu *et al*. (14) we calculated *B*^*R*^ = 3.2 × 10^−19^J from an isolated patch of our lipid membrane (see Supplementary Material S3). Taking the value *µ*_*s*_ = 11µNm/s from our *D*^*M*^ = 125 cell, this indicates the bending rigidity of the bilayer to incur a negligible effect on the whole-cell deformation response compared to the shear modulus (7). In a healthy RBC, the bending rigidity of the lipid bilayer is 2-3 orders of magnitude larger than the cytoskeleton (8). However, our model cytoskeleton is very simple in construction, being comprised of CG particles of uniform mass held together by standard 12-6 LJ potentials. Therefore, it is possible that the stretching response is being partly dampened by an uncharacteristically high bending rigidity in the cytoskeleton.

More generally, it is challenging to create direct comparison between the force-extension curves generated in this *in silico* study and past *in vitro* studies. Firstly, the force calibration methods of *in vitro* studies introduce large uncertainties, resulting in a wide range of reported force measurements (21, 23, 31). Mills *et al*. (21) presented major revisions to the force calibration of their initial work (22, 31), having initially estimated the maximum applied force to be double that from their revised calibration. Furthermore, how the axial diameter is defined can be contentious between *in vitro* experimentation and our CGMD simulations. In an optical tweezers experiment, the microbeads are physically bound to the cell surface. The axial extension *λ*_*x*_ can then be directly determined by measurement of the relative displacement of the two beads (21, 23). However, in our simulations we assume the effect from the microbeads by applying forces directly to designated regions of the lipid membrane. Without the presence of explicit beads, *λ*_*x*_ must be determined from the relative displacement of some part of the cell surface, made more challenging by the diffusive nature of the lipid particles. Following the recommendation of Siguenza *et al*. (23), we have defined *D*_*x*_ as the distance between planes intersecting perpendicular to the innermost “bead” lipids (see Figure 2B). This may have led to the under-representation of *λ*_*x*_ relative to the experiments of Mills *et al*. (21) and Suresh *et al*. (30) in the high force regime.

Another notable divergence with past studies occurs above the low-force regime. Previous studies have consistently shown a plateau in extension with increasingly large force, with an asymptote around *λ*_*x*_ ∼ 2 (12, 21–23, 30). This is not seen in our model. Instead, the rate of growth of *λ*_*x*_ (and decay of *λ*_*y*_) continues to increase with *F*_*x*_, until a critical point at which the membrane ruptures. Human RBC lysis due to critically high shear strain has been confirmed by many past *in vitro* studies (33–35). While RBCs are able to withstand linear extensions up to 250% (23), they are very susceptible to rupture from increasing surface area. Micro-pipette measurements showed rupture above a critical area strain of 2-4% (33), while cell-poking found rapid cell lysis above 2.6% area strain (35). This is comparable to when stretching our CGMD cells, with the *D*^*M*^ = 50 cell bursting at an area strain of 6.1%. Conversely, the cell sizes that reached the maximum applied force of *F*_*x*_ = 210pN without rupture had maximum area strain of order ∼ 1%. Past application of the Fu *et al*. model also found rupture to occur during the shape evolution phase, and when subject to high shear force from external fluid flow (18). Membrane rupture during the shape evolution phase was common in our simulations, hence the employed modifications to the mass and Yuan potential parameterisation to strengthen the particle bindings during this phase.

Rupture of the membrane is also conceptually reasonable in a CGMD model. Being a particle model operating through pair potentials, all attracting particles are bound within energy wells of finite depth. Introducing an external force to particular particles will add energy, acting towards breaking them free of these wells. The collection of CG lipids within the sphere of influence of each bead is kept constant, but have an evolving collective shape (see Figure 5). Initially, when the *XY* -plane of the cell is circular, these particles will be tightly spread within a circular patch on each edge of the membrane. However, as the cell is stretched along *x*, these regions are pulled outwards, becoming increasingly elongated. The Yuan potential binds the lipids together with a LJ-like relationship of inter-particle separation, within a cutoff radius *r*_*c*_. Therefore, the more particles within *r*_*c*_ of a particular CG lipid, the more tightly that particle will be bound to the whole cell. As each region of assigned bead particles is pulled into an increasingly elongated bulb, those lipids at the extremity will have fewer surrounding particles, thus lose binding strength to the whole cell. Eventually, the external force overcomes the restoring force of the summed inter-particle bindings and these lipid particles break free. This causes the whole cell to then lose structural integrity, and “pop”.

**Figure 5:**
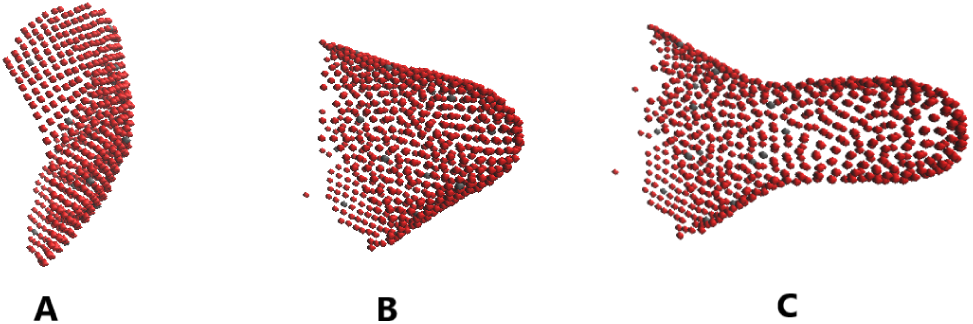
Graphic showing the shape evolution of a region of a cell stretched in the positive *x* direction, where only those *N*_+_ lipids designated bead particles are shown, being the only particles directly subjected to the external force. Before any stretching has occurred the cell is in its initial stable state, and the region of bead particles is relatively flat (**A**). At a late stage the cell has undergone a significant deformation, and the region has been pulled into a rounded cone (**B**). Finally, the extraneous particles start losing their binding with the whole cell and rupture becomes imminent (**C**).

Critically, it is also this force balancing between the internal restorative and external stretching forces that requires introduction of the force scaling term r_*F*_. Larger cells are comprised of more particles, but the proportional size of the optical microbeads is kept constant. Therefore, if the total force is kept constant, larger cells will have *F*_*x*_ spread over a larger number of bilayer particles. By maintaining the same model parameterisation between cell sizes, a cell comprising more particles will also have a greater sum of restoring forces within it, from the greater number of inter-particle potentials. Therefore, to produce the same extension in a larger cell as a smaller cell will require a larger total external force, proportional to the ratio in surface area. This leads to the 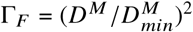 scaling term, which acts to maintain a constant force *per-particle*, rather than *total* force, as dictated by the model parameterisation.

A final consideration to the shape and scale of the force-extension curves is the model time-scale. Comparative RBC optical tweezer stretching experiments typically transition from zero to maximum strain over an order of 2-5s (22). The simulations performed in this work reached maximum strain after ≲ 1s. The rate of stretching is able to effect the temperature of a system *in vitro*, and thus the elastic properties of the components (22). The critical shear strain for RBC lysis has also been shown to be inversely proportional to the length of exposure to shear stress (36). Our fast deformation rate was due to having required a very short time-step size *τ* = 1.6ns, such to maintain numerical stability in the simulations. To then achieve maximum deformation on the order of 5 seconds would have required over 6 × 10^8^ time-steps, which for the *D*^*M*^ = 150 cell would expect to take up to 40 days to simulate.

## 5 CONCLUSION

The scale invariance of a CGMD model for a single RBC has been tested, first qualitatively through the shape evolution, then quantitatively by determination of the shear modulus *µ*_*s*_ through simulation of optical tweezer stretching. By evolving cells of various diameter from their initially spherical configurations, cells of size *D*^*M*^ ≥ 100 were found able to develop the characteristic biconcave discocyte shape of a healthy RBC. All evolved cells were then subjected to optical tweezer-style stretching, with *µ*_*s*_ in the range 6 − 15µN/m calculated from the low-force limit of resulting force-extension curves. Despite implementation of a dimensional scaling, a positive correlation was found between cell size and the deformation response over the range of *D*^*M*^ tested. This indicated an order of magnitude difference in measured shear modulus between a full-scale cell and one at 1/32 scale, thus contradicting a total scale invariance in the model. However, without being able to test cells of *D*^*M*^ *>* 200, the persistence of this trend is speculative. Furthermore, the similarities of the *D*^*M*^ = 100 − 200 miniature cells to a full-scale human RBC in shape and shear modulus indicate them to be valuable as simplified representations. Therefore, we determine the scaled use of miniature CGMD cells of *D*^*M*^ = 100 − 200 to be a valid approximation for the purposes of estimating the elastic responses of a cell in-silico. This finding supports the use of the miniature cell approach in further studies, with its considerable computational advantages opening up numerous possibilities in simulations of physically larger, more numerous and more complex CGMD cellular systems than have been performed to-date.

## AUTHOR CONTRIBUTIONS

P.A. carried out all research, simulations, data analysis and writing of the article. S.H. and A.M.S. designed the research, as the primary and secondary PhD supervisors.

## ACKNOWLEDGMENTS

We thank S.-P. Fu for their help in initial implementation of the model. This work was carried out using the computational facilities of the Advanced Computing Research Centre, University of Bristol - http://www.bris.ac.uk/acrc/.

An online supplement to this article can be found by visiting BJ Online at http://www.biophysj.org.

